# Correlated evolution of sex allocation and mating system in wrasses and parrotfishes

**DOI:** 10.1101/665638

**Authors:** Jennifer R. Hodge, Francesco Santini, Peter C. Wainwright

## Abstract

In accordance with predictions of the size-advantage model, comparative evidence confirms that protogynous sex change is lost when mating behavior is characterized by weak size advantage. However, we lack comparative evidence supporting the adaptive significance of sex change. Specifically, it remains unclear whether increasing male size advantage induces transitions to protogynous sex change across species, as it can within species. We show that in wrasses and parrotfishes (Labridae), the evolution of protogynous sex change is correlated with polygynous mating, and that the degree of male size advantage expressed by polygynous species influences transitions between different types of protogynous sex change. Phylogenetic reconstructions reveal strikingly similar patterns of sex allocation and mating system evolution with comparable lability. Despite the plasticity of sex determination mechanisms in labrids, transitions trend towards monandry (all males derived from sex-changed females), with all observed losses of protogyny accounted for by shifts in the timing of sex change to prematuration. Likewise, transitions in mating system trend from the ancestral condition of lek-like polygyny toward greater male size advantage, characteristic of haremic polygyny. The results of our comparative analyses are among the first to confirm the adaptive significance of sex change as described by the size-advantage model.

## Introduction

Sequential hermaphroditism is a reproductive strategy with multiple evolutionary origins distributed sporadically across the tree of life (Policansky 1982; Sadovy de Mitcheson and Liu 2008). It is characterized by a change in the functional expression of sex, from one to the other. Among vertebrates, sequential hermaphroditism is found only in teleosts (Todd et al. 2016), where sex change can be male-to-female (protandry), female-to-male (protogyny), or serial bidirectional. Each form of sequential hermaphroditism has evolved multiple times within teleosts, demonstrating the lability of fish sex determination mechanisms (Smith 1975; Charnov 1982; Policansky 1982; Mank et al. 2006).

The dominant theory describing the adaptive significance of sequential hermaphroditism is the size-advantage model (SAM) (Ghiselin 1969; Warner 1975; Leigh et al. 1976; Charnov 1982). The model contends that sex change is favored when the rate of increase in reproductive value with size and age differs between the sexes. Correspondingly, gonochorism (the existence of separate, fixed sexes) is predicted when size-specific male and female reproductive outcomes do not differ (Warner 1975; Muñoz and Warner 2003, 2004). A range of complex factors capable of contributing to differences in reproductive value between the sexes have been integrated into the SAM (Charnov 1982; Warner 1988). Of these, mating behavior has emerged as an important determinant of size-related differential reproductive outcomes (Shapiro 1987; Ross 1990; Munday et al. 2006a). This is because certain mating systems are also contingent on male size advantage.

For example, protogyny, the most prevalent form of sequential hermaphroditism in fishes (Sadovy de Mitcheson and Liu 2008; Todd et al. 2016), is predicted to be adaptive when reproductive value increases with size faster in males than in females (Warner 1975; Leigh et al. 1976). Protogynous species can be either monandric, in which case all males are derived from sex-changed females, or diandric where males are either born into the population or derived from sex-changed females (see table 1 for definitions). Sex-based size asymmetry is also characteristic of polygynous mating, where males use their size advantage to monopolize access to females by guarding them or the resources on which they depend (Ghiselin 1969; Taborsky 1998). This behavior results in large males having a reproductive advantage over females and small males, thereby enabling selection for protogynous sex change (Warner 1975, 1984, 1988). Polygynous mating can include haremic systems, where a single male monopolizes and mates with one (Pitcher 1992) or more females within a defined, permanent territory; and lek-like systems, where males establish temporary territories that are visited by females for the purposes of reproduction. As mating becomes more promiscuous, sperm competition increases and the reproductive advantage of large males decreases due to the dilution of gametes by other males, consequently reducing selection for protogyny (Warner 1975).

**Table 1:** Definition of key terms.

|  | Definition |
| --- | --- |
| Sex allocation: |  |
| Gonochorism | Individuals reproduce exclusively as either male or female throughout their lives. |
| Diandric protogyny | Males are either born into the population (primary males) or derived from sex-changed females (secondary males). |
| Monandric protogyny | All males are derived via sex change from functional females (secondary males). |
| Mating system: |  |
| Promiscuity | Terminal phase males do not defend territories with the purpose of attracting mates. |
| Lek-like polygyny | Terminal phase males establish temporary territories visited by females for the purposes of reproduction. |
| Haremic polygyny | Terminal phase males monopolize one or more females within a permanent territory. |

Sources. – Warner and Robertson 1978; Colin and Bell 1991; Sadovy de Mitcheson and Liu 2008.

When large males have strong social control over females, as in haremic systems, monandric protogyny is predicted (Robertson and Choat 1974; Robertson and Warner 1978; Warner and Robertson 1978; Warner 1984; Nemtzov 1985). In lek-like systems where the social control of large males is reduced, primary males are able to realize reproductive success and selection should favor diandric protogyny (Robertson and Choat 1974; Emlen and Oring 1977; Robertson and Warner 1978; Warner and Robertson 1978). Correspondingly, gonochorism is predicted when males lose social control as in promiscuous mating behaviors such as group spawning (Robertson and Warner 1978; Warner 1984; Hoffman 1985). Qualitative assessments of mating behavior support its role as a primary determinant of the degree of size advantage, and consequently of the incidence and direction of sequential hermaphroditism in fishes (Warner 1984; Ross 1990; Munday et al. 2006a; Erisman et al. 2013).

Population demographic studies provide empirical support for the effect of mating behavior on sex allocation within single species, where differences in population size and the distribution of resources affect the ability of large males to maintain permanent territories, thereby inducing predicted changes in sex allocation (Warner and Robertson 1978; Warner and Hoffman 1980a,b). Phylogenetic comparative studies have shown that transitions from protogyny to gonochorism are contingent upon weaker size advantage (Kazancıoğlu and Alonzo 2010) and group spawning (Erisman et al. 2009). However, no comparative studies have supported predictions about the adaptive significance of protogynous sex change. Specifically, we lack comparative evidence showing that as size-advantage increases and large males have greater opportunity to monopolize mating, protogynous sex change evolves. Many aspects of an organism’s biology, demography and ecology have the potential to affect the expression of complex traits such as mating system and sex allocation. Moreover, behavioral traits tend to be more evolutionarily labile than life-history traits (Blomberg et al. 2003). It remains unknown whether variation in the degree of male size advantage characteristic of specific types of polygynous mating induces evolutionary transitions to and within types of protogynous sex change in the context of other influential factors. Do the effects of mating behavior on sex allocation observed within species scale up to macroevolutionary patterns?

The wrasses and parrotfishes, along with cales and weed-whitings (Labridae) provide an ideal opportunity to evaluate the evolutionary synergy between sex allocation and mating behavior along a spectrum of male size advantage. The Labridae form a monophyletic assemblage (Aiello et al. 2017) comprising one of the largest families of marine fishes with a global distribution spanning tropical and temperate waters. Extensive scientific interest in labrid mating and sex systems has produced some of the most influential insights into the adaptive significance of sequential hermaphroditism (Darwin 1871; Ghiselin 1969; Robertson 1972; Robertson and Choat 1974; Warner 1975; Warner et al. 1975; Leigh et al. 1976; Muñoz and Warner 2004), as well as many observations that both support and contradict predictions of the SAM (Robertson and Warner 1978; Warner 1984; Nemtzov 1985; Warner and Lejeune 1985; Cowen 1990; Morrey et al. 2002; Adreani et al. 2004; McBride and Johnson 2007). The most comprehensive comparative analysis of the SAM to date focused on labrids, whereby the authors combined mating behavior with other phenotypic traits, including colour, to quantify male size advantage as either strong or weak (Kazancıoğlu and Alonzo 2010). They found that protogyny is less likely to be lost under strong size advantage but did not find evidence that strong size advantage induces transitions from gonochorism to protogyny. Incorporating variation within polygyny and protogyny will provide more detail about the evolutionary dynamics between mating behavior and sex allocation, and allow us to assess the effects of each trait regime on the adaptive evolution of the other.

As a result of considerable past research efforts, labrid mating and sex systems are comparatively well quantified. Protogynous sex change is pervasive among labrids and has been reconstructed as the ancestral condition (Sadovy de Mitcheson and Liu 2008; Kazancıoğlu and Alonzo 2010; Erisman et al. 2013). Labrid species express both types of protogyny – monandry and diandry – the origins of which have yet to be infered in the context of a time-calibrated phylogeny. The family also includes some gonochoristic species. Mating systems are equally as diverse; some species maintain harems, other species exhibit lek-like polygyny, and other species mate promiscuously with no territory defense by males for the purpose of attracting mates. Finally, robust phylogenetic hypotheses exist that include over half of the nominal labrid species, with opportunities to expand the taxonomic representation of species arising frequently.

We are now in a position to explore the evolutionary history of, and synergy between, mating behavior and sex allocation in wrasses and parrotfishes along a continuum of male size advantage. We use Bayesian methods to reconstruct the evolutionary history of each trait in the context of a new, taxonomically-expanded phylogeny and apply discrete trait comparative methods to test predicted associations between specific types of protogynous sex change and polygynous mating with distinct degrees of male size advantage.

## Materials and Methods

### PHYLOGENETIC INFERENCE AND DIVERGENCE TIME ESTIMATION

We reconstructed phylogenetic relationships using a molecular dataset comprised of four mitochondrial (12S, 16S, COI, and CytB) and three nuclear loci (RAG2, TMO4c4, and S7), with a total of 4,578 base pairs. Sequence data were compiled from GenBank for all available nominal species – at the time of sampling this included 403 species from 74 genera, and two outgroup taxa (see table A1 for accession numbers and table A2 for information on molecular sampling). Gene sequences were aligned separately in Geneious Pro 8.1.9 (Biomatters, Auckland, New Zealand; http://www.geneious.com/) using default settings and the alignments were manually adjusted through the insertion or deletion of gaps and trimmed to minimize the amount of missing data. Alignments were concatenated and partitioned by gene region, with separate partitions for the 3^rd^ codon position of protein coding genes.

We performed Bayesian inference (BI) using partitioned mixed models in MrBayes (Ronquist and Huelsenbeck 2003) to estimate the tree topology and branch lengths. We sampled across the entire general time reversible (GTR) model space using reversible jump Markov chain Monte Carlo (rjMCMC) methods to integrate out uncertainty about the correct substitution model for each partition (nst = mixed). This allowed us to quantify posterior probabilities of the substitution models sampled (table A3). Parameters were unlinked across partitions. Substitution rates and stationary nucleotide frequencies were allowed to evolve under different rates using a flat Dirichlet prior. The shape of the gamma distribution of rate variation evolved under an exponential prior with a mean of one. Branch lengths were unconstrained under an exponential prior with a rate of 200 (mean = 0.005). Posterior probabilities of clades were calculated following two 40 million generation Markov chain Monte Carlo (MCMC) analyses, each with eight chains (temp = 0.02) and two swaps, sampling every 2,000 generations. Convergence was assessed in Tracer (Rambaut et al. 2009). Upon examination of the trace files, a conservative burn-in of 45% was discarded from each run and a 50% majority-rule consensus tree was computed using the remaining sampled trees.

The majority-rule consensus tree was converted to a rooted, ultrametric tree using the chronos function in the R package ape (Paradis et al. 2004). We set the smoothing parameter lambda to 0.9 and specified a relaxed model of substitution rate variation among branches and seven age constraints (table A4). We performed a divergence time analysis in BEAST (Drummond et al. 2012) using the resultant ultrametric tree as the starting tree. Partitioning followed the scheme above and models of molecular evolution were specified using the parameters and priors corresponding to the model with the highest posterior probability from the MrBayes analysis and empirical base frequencies (see table A3 for substitution model details). Divergence times were estimated under a relaxed uncorrelated lognormal clock model (Drummond et al. 2006) and the birth-death process (Gernhard 2008). Evidence from six fossils informed exponential prior distributions on corresponding nodes (table A4) to time-calibrate the trees. Monophyly of the Labridae was enforced. Posterior samples from three independent MCMC analyses, each with 80 million generations, sampling every 4,000 generations, were assessed for convergence and appropriate burn-in in Tracer (Rambaut et al. 2009). Tree files were combined using LogCombiner (Drummond et al. 2012) following the removal of 27.5–40% burn-in, and resampling every 16,000 states (deposited in the Dryad Digital Repository: URL pending publication; Hodge et al. 2019). A maximum clade credibility tree was constructed using TreeAnnotator (Drummond et al. 2012) to display median ages and 95% highest posterior density (HPD) intervals (fig. 1; deposited in the Dryad Digital Repository: URL pending publication; Hodge et al. 2019).

**Figure 1:**
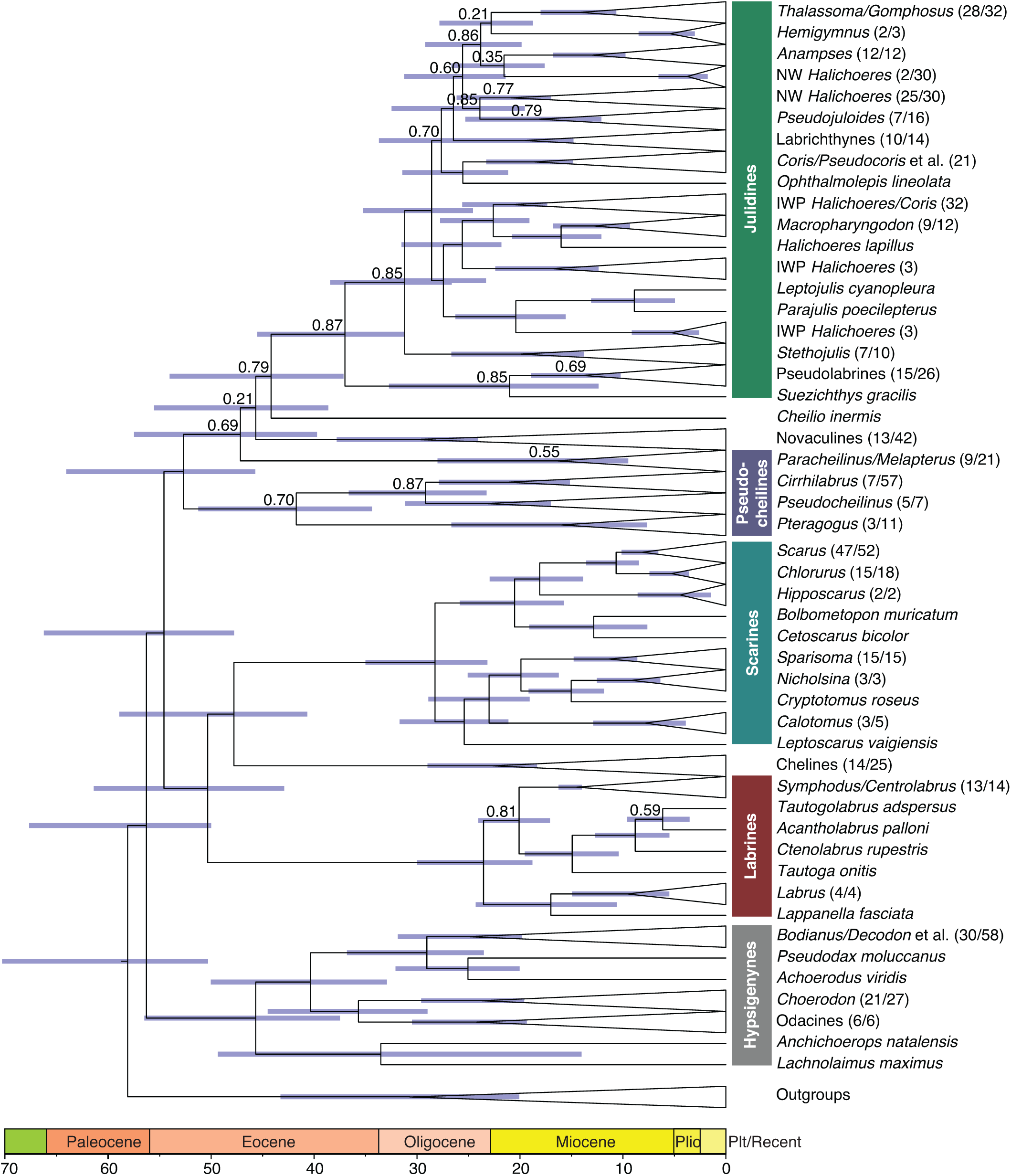
The maximum clade credibility tree estimated using Bayesian inference shows relationships between major clades of wrasses and parrotfishes and median node ages in millions of years before present, with 95% HPD intervals denoted by bars at nodes. Posterior probability support for nodes was ≥ 0.9 unless otherwise indicated. Tree files available online in the Dryad Digital Repository: URL pending publication (Hodge et al. 2019).

### TRAIT DATA COMPILATION

We compiled data on species-specific mating systems and sex allocation pathways from the primary literature (tables 1, A5; deposited in the Dryad Digital Repository: URL pending publication; Hodge et al. 2019). Mating system classifications focused only on terminal phase males and did not consider the mating strategies of initial phase males – although it is known that the reproductive output of initial phase males can outweigh that of terminal phase males for some species dependent on location-specific population dynamics (Warner and Hoffman 1980a, 1980b; Warner 1982). We applied the consensus classification of the predominant mating system (i.e. supported by multiple authors) whenever possible, and otherwise relied on the most recent observations. We restricted sexual ontogeny data to accounts of protogyny that were distinguishable as either monandric or diandric based on gonad histology, population demographics or both. Cases where males are derived from females that have not passed through a functional stage were categorized as functionally gonochoristic following previous work (Sadovy and Shapiro 1987; Sadovy de Mitcheson and Liu 2008; Kazancıoğlu and Alonzo 2010; Erisman et al. 2013). Mating system and sex-change data were available for 89 labrid species (table A5).

### ANCESTRAL STATE RECONSTRUCTION

We reconstructed the evolutionary history of sex allocation and mating system using the MultiState package implemented in BayesTraits (Pagel et al. 2004; Pagel and Meade 2006). We fit continuous-time Markov models to each set of discrete character data using a rjMCMC analysis to derive posterior distributions of the ancestral state and transition rates. An exponential reversible jump hyperprior (0 10) was specified for the rate parameter distributions, and the trees were scaled to have a mean branch length of 0.1. Markov chains were run three times across a random sample of 1,000 time-calibrated phylogenies (deposited in the Dryad Digital Repository: URL pending publication; Hodge et al. 2019) for four-million iterations, sampling every 4,000 steps, following a burn-in of one-million iterations. We monitored the average acceptance rates to ensure the values were between 20 and 40%, indicating that the rjMCMC was mixing well. We examined traces of the likelihood and parameters in Tracer (Rambaut et al. 2009) to ensure convergence and effective sample sizes (ESS > 200) across the three independent runs. Parameter summary statistics were calculated from the concatenated estimates of three converged runs.

To visualize the evolutionary history of each trait we also performed ancestral state reconstructions as described above on the MCC tree and calculated the average posterior probabilities of each character state at each node in the phylogeny (fig. 2). Transitions in character states were defined as nodes with posterior probability values ≥ 0.50 in support of a transition relative to a preceding node (i.e. a direct ancestor) with posterior probability ≥ 0.50 for a different state and included changes along terminal branches. We summarized the number, location and nature of these transitions.

**Figure 2:**
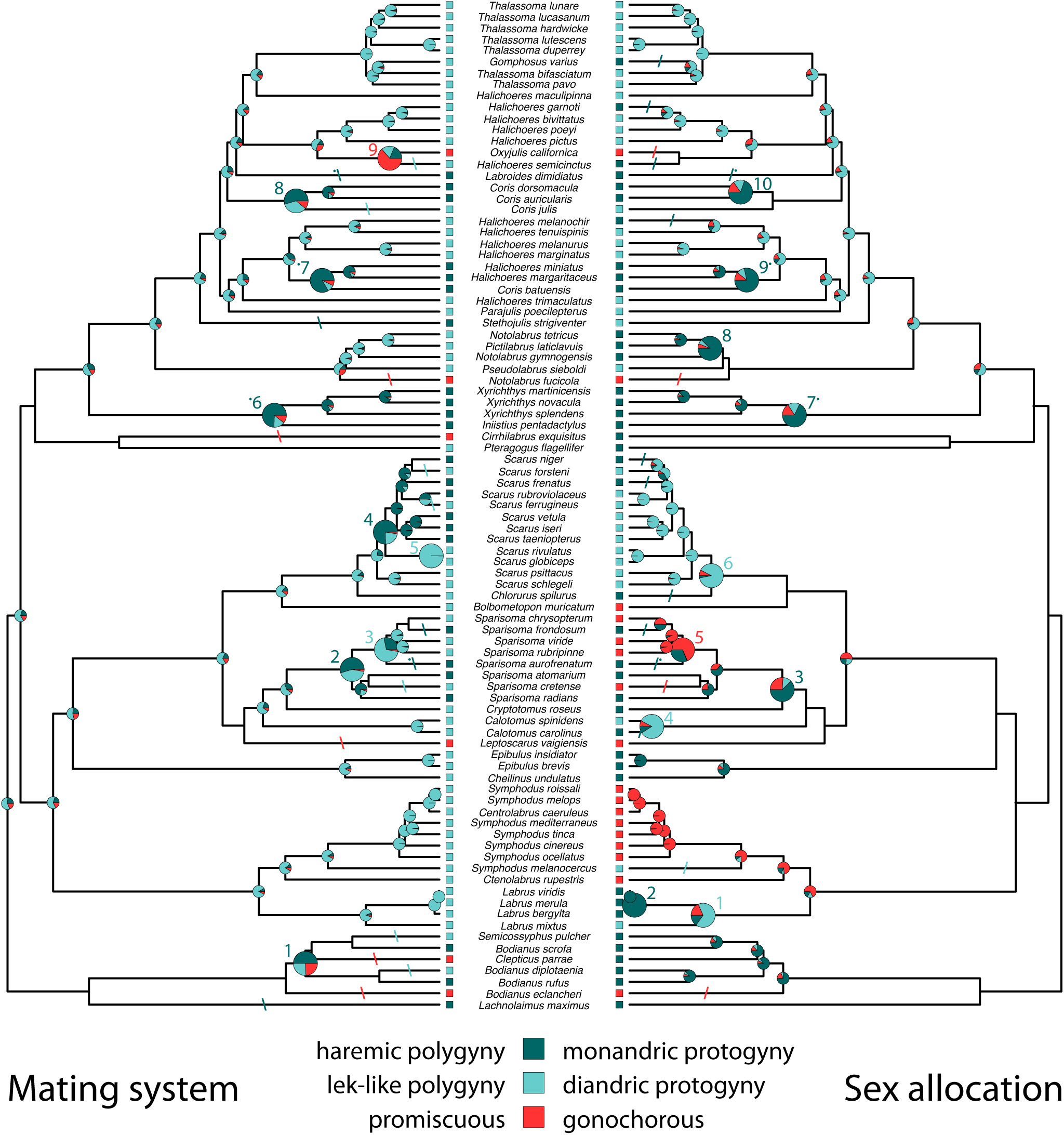
Bayesian ancestral reconstructions of the evolutionary history of mating system and sex allocation. Character state colors correspond to predicted associations. Pie charts at nodes are only shown for ancestral states resolved with ≥ 0.50 posterior support. Enlarged pie charts and corresponding numbers indicate evolutionary transitions, defined as nodes with posterior probability values ≥ 0.50 in support of a transition relative to a preceding node with posterior probability ≥ 0.50 for a different state. Evolutionary transitions along terminal branches are indicated by dashes at the mid-point of the terminal branch. In the interest of visual clarity these transitions are not numbered but can be counted from bottom to top. Dots next to numbers and dashes indicate simultaneous transitions to states with predicted associations. Data and tree files available online in the Dryad Digital Repository: URL pending publication (Hodge et al. 2019).

### TRAIT CORRELATIONS

To test whether increasing male size advantage results in transitions to protogynous sex change we compared the fit of independent and dependent models of trait evolution using the Discrete package implemented in BayesTraits (Pagel et al. 2004; Pagel and Meade 2006). The independent, or null, model of evolution assumes there is no correlation between two traits and that they evolved independently. The dependent model describes the correlated evolution of two traits such that the rate of change in one trait depends on the state of the other trait. As Discrete accepts only binary trait data we performed two tests: the first used the full dataset (n = 89 species) to assess the correlation between polygyny and protogyny (species coded as either promiscuous or polygynous and gonochorous or protogynous); the second used a reduced dataset that included only species that are both polygynous and protogynous (n = 70 species) to assess predicted correlations between the two traits based on different degrees of male size advantage (species coded as either lek-like or haremic and diandric or monandric).

Models were fit using the same set of 1,000 trees, number of generations, sampling frequency, burn-in and hyperprior specifications as the MultiState analysis above. Run diagnostics of the acceptance rates, likelihood and parameter traces were also performed as above. We determined the most probable evolutionary model by calculating log Bayes Factors (BF) for each pair of models as twice the difference in the log marginal likelihood of the dependent model minus the independent model (Kass and Raftery 1995). Marginal likelihoods were estimated using the stepping stone sampler (Xie et al. 2011) implemented in BayesTraits (Pagel et al. 2004; Pagel and Meade 2006), where each independent run sampled 100 stones, each with 10,000 iterations. Log BFs were averaged across independent runs. The log BF quantifies the weight of evidence against the null hypothesis (the independent model) whereby values less than two indicate little evidence, values from two to five indicate positive evidence and values greater than five indicate strong evidence for the dependent model over the independent model (Raftery 1996). We calculated Z-scores for each transition parameter as the proportion of transitions assigned to zero across the three independent, concatenated runs. The Z-score provides an additional descriptor of the likelihood distribution of the transition rate. Low Z-scores indicate that transitions were rarely assigned to zero and are likely to occur, whereas Z-scores close to one describe transitions that were frequently assigned to zero, indicating that they are unlikely to occur.

## Results

### PHYLOGENETIC INFERENCE AND DIVERGENCE TIME ESTIMATION

Our expanded taxonomic sampling represents 60.15% of nominal species and 86.05% of nominal genera (table A6). Our analysis included representative species from all but 12 genera, eight of which are monotypic (table A6). Molecular data for all nominal species were obtained for 35 genera (40.7% of all genera), and sixty-two of the 74 genera included in our analysis (83.8%) had molecular data available for over half of their nominal species. Twelve genera had molecular data for less than half of their nominal species, including the second largest genus *Cirrhilabrus*. Molecular data coverage for the species in our dataset ranged from 31.76% of species with sequence data (S7) to 81.98% (16S) of species with sequence data (table A2). On average, molecular coverage was 1.7 times higher for mitochondrial than nuclear gene regions.

Our phylogenetic hypothesis, represented by the MCC tree, was well-resolved with high posterior probability support for most nodes (fig. 1). Log files from three independent BEAST analyses showed high effective sample sizes for most parameters (posterior ESS values > 200 for three combined analyses), indicating valid estimates based on independent samples from the posterior distribution of the Markov chain. The MCC tree was compiled from 9,625 post-burn-in trees (38,500,000 generations).

Strong posterior probability support was obtained for the deepest nodes of the phylogeny, but considerable uncertainty emerged for the nodes leading into the julidines (specifically at the splits between the pseudocheilines, the novaculines and the julidines), where posterior probability support for the relationships between genera decreases (fig. 1). These three clades comprise the largest tribe (Julidini) and the least well sampled tribes (Chirrhilabrini and Novaculini), thus future studies should prioritize the interrelationships of species in these groups.

We recovered moderate to high posterior probabilities (between 0.79 and 1) supporting the monophyly of 25 genera (48.08% of all non-monotypic genera; table A6). Five genera were recovered as paraphyletic (posterior probabilities 0.37–1), and eight genera were recovered as polyphyletic, (posterior probabilities 0.33–1). Monophyly remains undeterminable for 14 non-monotypic genera due to lack of species sampling (26.92% of all non-monotypic genera). Of these 14 genera, 10 were represented in the analysis by only one species.

We estimated the origin of the Labridae to be in the late Paleocene around 56.28 Ma (95% HPD: 67.15–50), nearly 10 Myr after the Cretaceous–Paleogene (K–Pg) extinction event (Renne et al. 2013) and 0.8 Myr prior to the Paleocene-Eocene Thermal Maximum (McInerney and Wing 2011). Subsequent divergence within the family began approximately 54.3 Ma (95% HPD: 66.05–48.3) and continued throughout and beyond the early Eocene climatic optimum 52.6–50.3 Ma (Payros et al. 2015). All non-monotypic genera were established during the late Oligocene through the Pleistocene, between 23.66 and 0.39 Ma (95% HPD: 28.8–0.16). Most species diverged during the Pliocene and Pleistocene between 5.25 and 0.05 Ma (median estimated divergence of terminal taxa = 3.89 Ma).

### EVOLUTIONARY HISTORY OF SEX CHANGE AND MATING SYSTEM

Bayesian analyses indicated that the ancestral labrid mating system was most likely lek-like polygyny [average posterior probability = 0.57, 95% highest posterior density (HPD) interval: 0.3–1] but did not resolve the ancestral sex allocation pathway, as all three possible states had comparable posterior probabilities of ∼0.33 (fig. 2). Twenty-six transitions were recovered from Bayesian analyses of the MCC tree for both mating system and sex allocation. Nine of the 26 transitions in mating system occurred at the nodes and 17 along terminal branches. Transitions out of lek-like polygyny (61.5%) and to haremic polygyny (42.3%) were the most frequent. Ten of the 26 transitions in sex allocation occurred at the nodes and 16 along terminal branches. Transitions out of diandric protogyny (61.5%) and to monandric protogyny (65.4%) were the most frequent.

Transitions in mating system and sex allocation are tightly coupled. The effectively equal number of state-dependent transitions summarized on the MCC tree provide little resolution regarding the predominant effects of one trait regime over the other. Focusing on predicted character-state associations, we recovered seven state-dependent transitions in mating system (i.e. where mating system transitions to the predicted state given the state of sex allocation; nodes 1, 2 and 5; tip transitions 6, 10, 11 and 15) and eight in sex allocation (i.e. where sex allocation transitions to the predicted state given the mating system state; nodes 1, 4, 6 and 10; tip transitions 2, 8, 9 and 14), with another seven simultaneous transitions where mating system and sex allocation transition at the same node or along the same terminal branch (indicated by dots next to numbers and dashes in fig. 2).

From the Bayesian analyses of 1,000 tree topologies, transitions between lek-like and haremic polygyny had the highest rates and likelihoods (fig. 3, 5*A*). Promiscuity evolved with a higher rate and likelihood among lineages with haremic polygyny but, for haremic lineages, transitions to promiscuity were less likely than reversals to lek-like polygyny. For sex allocation, transitions to monandric protogyny had the highest rates and likelihoods (fig. 4, 5*B*). Transition rates between gonochorism and monandric protogyny are high in both directions, while transitions to and from diandric protogyny are unidirectional. Specifically, transitions to diandric protogyny rarely occur among monandric lineages and diandric lineages rarely transition to gonochorism.

**Figure 3:**
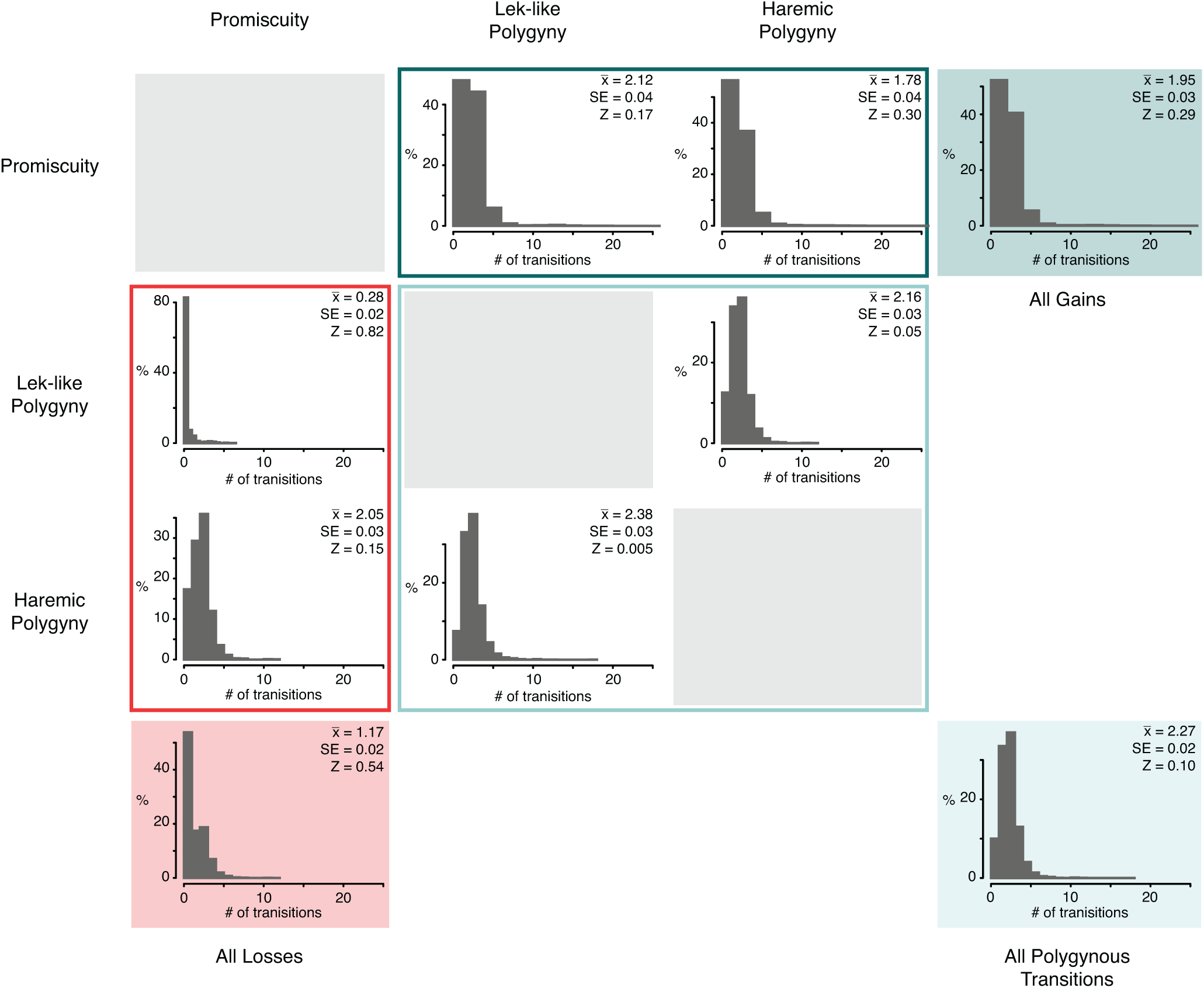
Posterior distributions of the rate coefficients characterizing transitions in mating system estimated from 1,000 Bayesian posterior distribution trees. Rows and columns indicate the pre- and post-transition states, respectively. Sample means (*x̅*), standard errors (SE) second, and Z-scores are provided for each transition. All transitions to polygynous mating are summarized in All Gains, all transitions to promiscuity are summarized in All Losses, and all transitions between lek-like and haremic polygyny are summarized in All Polygynous Transitions.

**Figure 4:**
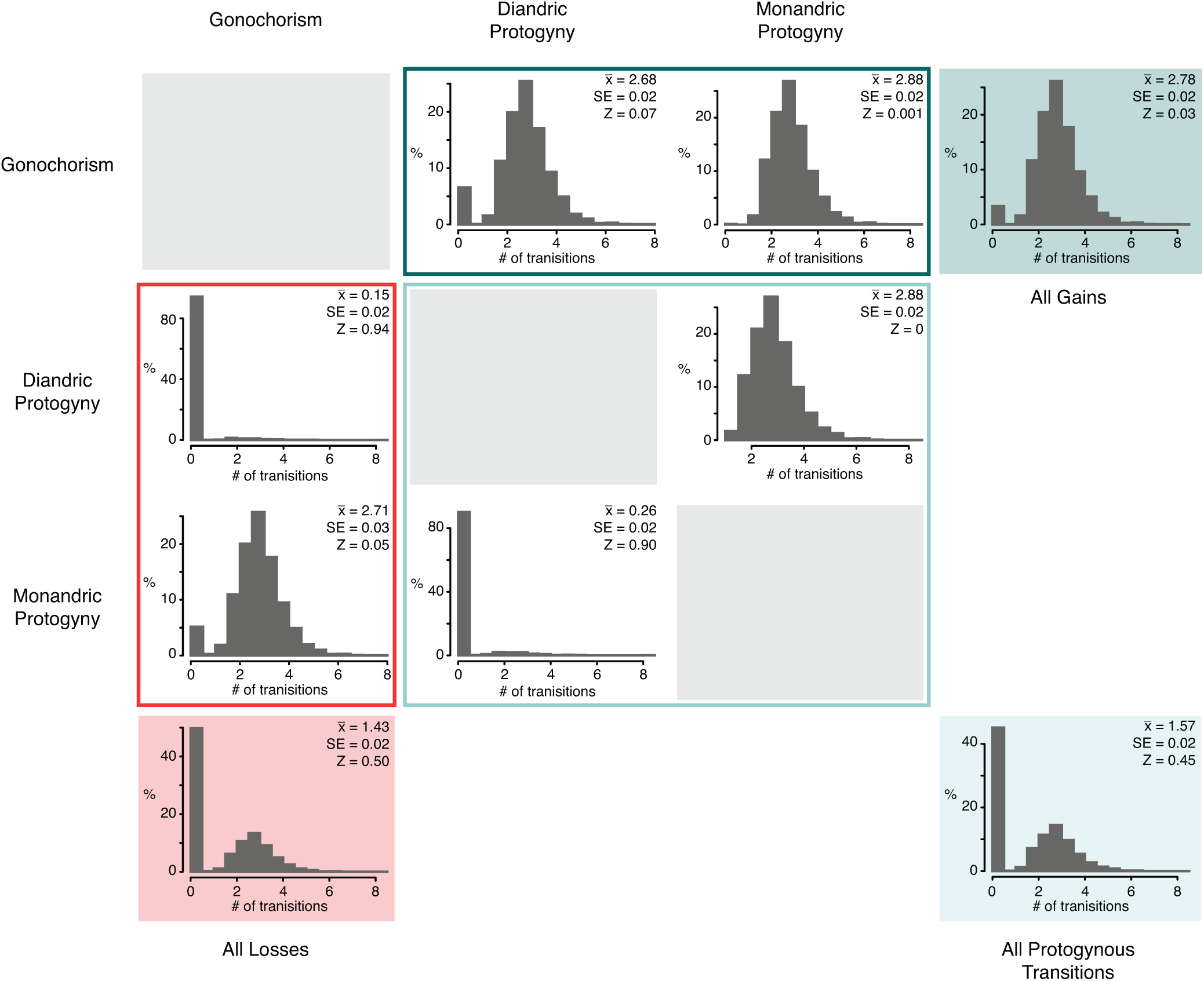
Posterior distributions of the rate coefficients characterizing transitions in sex allocation estimated from 1,000 Bayesian posterior distribution trees. Rows and columns indicate the pre- and post-transition states, respectively. Sample means (*x̅*), standard errors (SE) second, and Z-scores are provided for each transition. All transitions to protogynous sex change are summarized in All Gains, all transitions to gonochorism are summarized in All Losses, and all transitions between diandric and monandric protogyny are summarized in All Protogynous Transitions.

**Figure 5:**
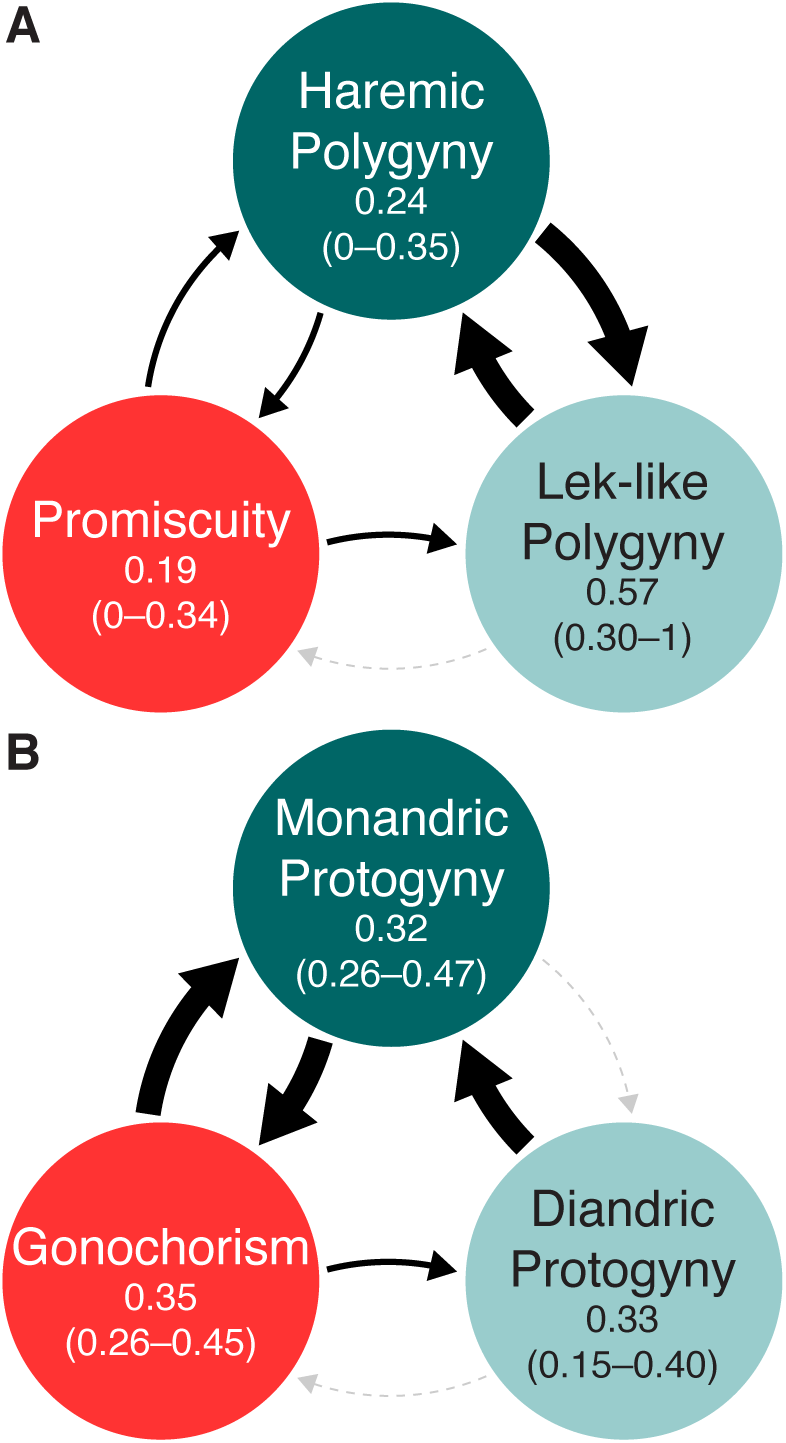
A visual representation of the likelihood of transitions between mating system (*A*) and sex allocation (*B*) character states. Relative transition probabilities are indicated by line weight, where thick solid lines represent Z-scores ≤ 0.05, thin solid lines represent 0.49 > Z > 0.06, and dashed lines represent Z-scores ≥ 0.5. Line colour indicates the rate class to which transitions were assigned most frequently, where rates assigned to ‘Z’ are shown in grey and rates assigned to ‘0’ are shown in black. Below each character state, the posterior probability of reconstructing that state as ancestral is shown (average and 95% highest posterior density interval).

### EVOLUTIONARY CORRELATIONS

Bayesian analyses show strong support for the correlated evolution of polygynous mating and protogynous sex change (average log Bayes Factor = 5.83; fig. 6*A* and table A7). Protogynous sex change is lost at a lower rate and with lower probability among polygynous lineages, than those that are promiscuous (91.6% of posterior samples had lower transition rates under polygynous mating). Polygynous lineages transition to protogynous sex change at a higher rate and with greater probability, than those that are promiscuous (51.0% of posterior samples had higher transition rates under polygynous mating). Transitions between promiscuous and polygynous mating showed some dependence on the state of sex allocation – specifically protogynous lineages transitioned to polygyny at a higher rate than gonochorous lineages, but only 41.2% of posterior samples reflected this state-dependent rate difference.

**Figure 6:**
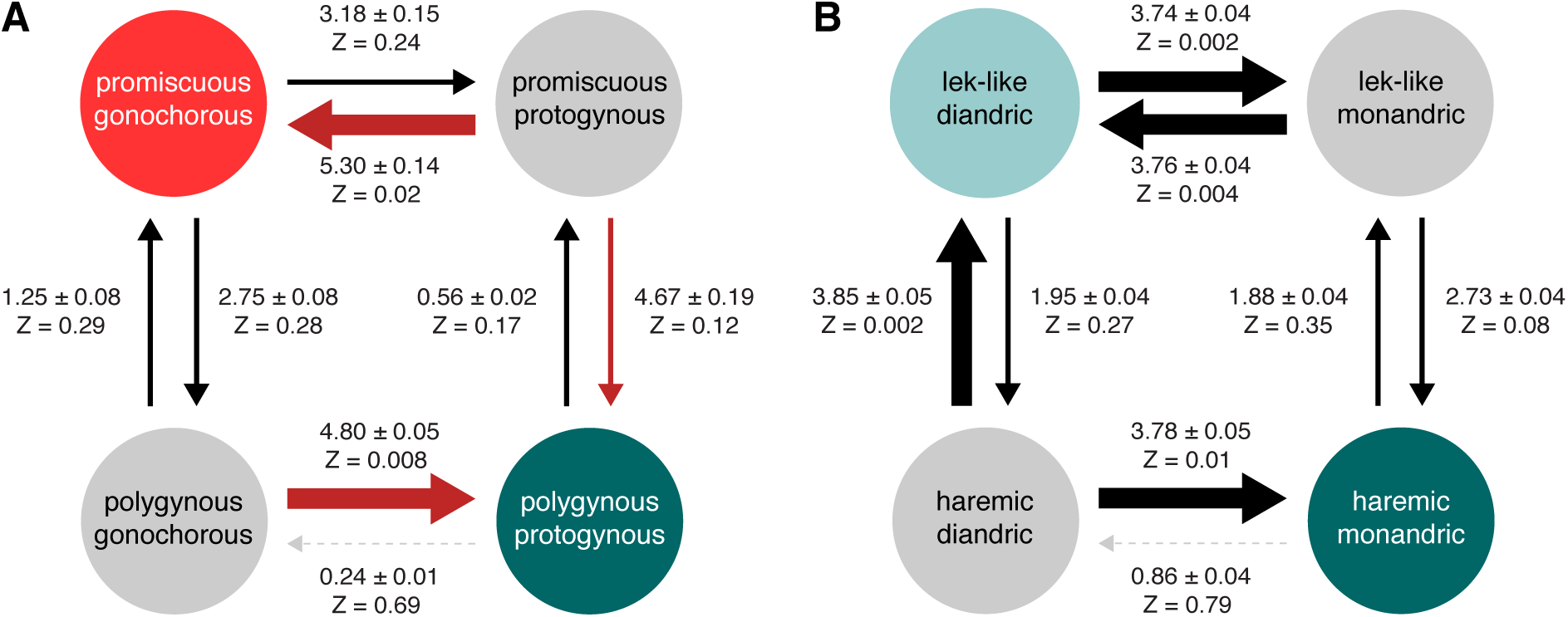
Mean evolutionary transition rates (*q*_xy_ ± SE) derived from Bayesian models of discrete character evolution fit to predicted associations between polygynous mating and protogynous sex change (*A*); and between specific types of polygynous mating and protogynous sex change (*B*). Below each transition rate, Z-scores denote the proportion of transition rates assigned to zero – values ≤ 0.05 are rarely assigned to zero and considered probable evolutionary events. Transition probabilities are also indicated by line weight, where thick solid lines represent Z-scores ≤ 0.05, thin solid lines represent 0.49 > Z > 0.06, and dashed lines represent Z-scores ≥ 0.5. Line colour indicates the rate class to which transitions were assigned most frequently, where rates assigned to ‘Z’ are shown in grey, rates assigned to ‘0’ are shown in black, and rates assigned to ‘1’ are shown in red.

Specific types of polygynous mating and protogynous sex change are also evolutionarily correlated as predicted by the SAM (average log Bayes Factor = 3.52; fig. 6*B* and table A7). Haremic lineages transition from monandric to diandric protogyny at a lower rate and with lower probability, than those with lek-like polygyny (80.9% of posterior samples had lower transition rates under haremic mating). However, transitions to monandry occurred at similar rates and with similar probability among lineages with either type of polygynous mating (94.8% of posterior samples had similar probabilities under each type of polygynous mating). Finally, the state of sex allocation does have some effect on the rate of transition between different types of polygynous mating, whereby diandric lineages revert to lek-like mating at a higher rate than monandric lineages, but only 42.8% of posterior samples reflect this state-dependent rate difference.

## Discussion

Evolutionary patterns of mating behavior and sex allocation across species of wrasses and parrotfishes are consistent with intraspecific patterns and predictions of the SAM. Our results confirm that protogynous sex change is less likely to be lost under polygynous mating where male-size advantage is stronger, and provide some of the first comparative evidence to support the evolution of protogynous sex change with increasing male size advantage. As male control over reproductive access to females increases from promiscuous to polygynous mating, so too does the size-dependent reproductive output of males relative to females, resulting in transitions from gonochorism to protogyny. We also find that specific types of polygynous mating and protogynous sex change have coevolved, following predictions of the SAM (Robertson and Choat 1974; Robertson and Warner 1978; Warner and Robertson 1978; Warner 1984). Our results support mating behavior as an important driver of transitions in sex allocation, with limited support for effects of sex allocation on mating behavior. The tight evolutionary coupling of the two trait regimes is likely facilitated by the labile nature of sex determination in fishes, allowing it to be less phylogenetically patterned than other life-history or physiological traits (Blomberg et al. 2003). Despite the lability of sex allocation, we found that monandric lineages rarely transition directly to diandric protogyny, instead appearing to transition through gonochorism on the pathway from monandry to diandry. We discuss how functional gonochorism has influenced this result and the overarching evolutionary trends of sex allocation in labrid fishes.

We have built upon previous efforts to reconstruct the evolutionary history of the Labridae (Westneat and Alfaro 2005; Alfaro et al. 2009; Cowman and Bellwood 2011; Santini et al. 2016; Aiello et al. 2017; Wainwright et al. 2018) and added 63 species to the phylogeny for a total of 403 out of 670 nominal species (as of March 20th, 2018; Eschmeyer et al. 2018). We note that the number of nominal species within Labridae has also increased by 8 since the last published phylogeny (Aiello et al. 2017). Topological relationships, including the non-monophyletic nature of thirteen genera (table A6), and divergence time estimates of the major groups that comprise the crown labrids were largely consistent with previous studies (Westneat and Alfaro 2005; Alfaro et al. 2009; Cowman and Bellwood 2011; Baliga and Law 2016; Aiello et al. 2017), as were estimates of extant species divergence (Hodge and Bellwood 2015). Our estimate of the origin of the Labridae in the late Paleocene (∼56.29 Ma; 95% HPD: 67.68–50) is on the younger end of recent estimates but within their 95% HPD intervals (Alfaro et al. 2009; Cowman and Bellwood 2011; Baliga and Law 2016; Aiello et al. 2017).

Lek-like polygyny can be traced back to the origin of the Labridae – as the estimated ancestral state with a high rate of reversal (fig. 2, 3, 5), it appears to be evolutionarily stable with the potential to affect detectable change in related traits. Labrid lineages have independently transitioned to haremic polygyny 11 times, each time transitioning from lek-like polygyny (fig. 2). Haremic polygyny is estimated to have arisen as early as 45.7 Ma (95% HPD: 55.5–38.6) along the lineage leading to the most recent common ancestor of the Novaculini (earlier transitions are also possible, specifically after the initial split of the Hypsigenyini [56.3 Ma; 95% HPD: 55.5–38.6] along the branch leading to *Lachnolaimus maximums*). In contrast, most transitions to promiscuous mating occur along much shallower, terminal branches, suggesting that, for labrids, this type of mating system may be less evolutionarily stable. Furthermore, Bayesian analyses show that promiscuous mating is more likely to arise from haremic polygyny than from lek-like polygyny (fig. 3, 5*A*), suggesting that promiscuity is a secondarily derived state. However, lineages that exhibit haremic polygyny are more likely to revert back to lek-like polygyny than they are to transition to promiscuity. This is concurrent with the expectation that differential dominance relationships between males will form when access to females cannot be controlled (Emlen and Oring 1977).

Protogyny was previously reported as the ancestral labrid condition (Sadovy de Mitcheson and Liu 2008; Kazancıoğlu and Alonzo 2010; Erisman et al. 2013). Here we distinguished between the different types of protogyny (monandric and diandric) but were not able to resolve the ancestral condition. Our result is based on the limited number of species for which sex allocation pathways are known, as this number increases so too may our ability to resolve the ancestral condition. The Novaculini and Julidini shared a diandric common ancestor approximately 45.7 Ma (95% HPD: 55.5–38.6) – the earliest sex allocation pathway reconstructed with confidence (fig. 2). Within this clade, the majority of transitions (82%) were to monandric protogyny and, more broadly, such transitions occurred with high rates (fig. 2, 4). Gonochoristic ancestors were reconstructed for other major clades including the Scarini and the Labrini (respective divergence time estimates: 28.3 Ma, 95% HPD: 35.0–23.2 and 23.5 Ma, 95% HPD: 30.0–18.8), while monandric protogyny emerged later in the evolutionary history of labrids, within the Hypsigenyini, Cheilinini and other clades (respective divergence time estimates: 20.0 Ma, 95% HPD: 25.0–16.0 and 12.3 Ma, 95% HPD: 16.8–8.4).

High transition rates between gonochorism and monandry (fig. 5*B*) lend further support to the ephemeral or non-existent nature of intermediate states in such transitions (Erisman et al. 2013) and show that diandry is not a necessary intermediate. Interestingly, transitions between gonochorism and monandry were largely restricted to the Sparisomatinae. All of the lineages that underwent transitions from monandry to gonochorism (with the exception of the lineage preceding *Leptoscarus vaigiensis*) gave rise to extant species that are functionally gonochoristic, with males derived from females that have not passed through a functional stage (Sadovy and Shapiro 1987; Sadovy de Mitcheson and Liu 2008). We note that simply because individuals are capable of prematurational sex change does not necessitate that all males be derived in this way. Some individuals could undergo postmaturational sex change, with the interval for sexual differentiation spanning pre- and postmaturation (Kazancıoğlu and Alonzo 2010). In this case, sex allocation would be more akin to diandric protogyny (Robertson et al. 1982; Munday et al. 2006b). Two features support the existence of ‘functional diandry’ in several of these species: smaller testes size of terminal phase males relative to initial phase males (Robertson and Warner 1978) – a common characteristic of other diandric species (Molloy et al. 2007), and polygynous lek-like mating (see further discussion below).

Regardless of the nature of these transitions, our results show that once lineages evolve the ability of some or all individuals to function first as females, they rarely lose it. Transitions trend away from diandry towards monandric protogyny, suggesting that when sustainable, labrids likely incur considerable fitness benefits by functioning first as females (or considerable fitness costs by not doing so). Labrids, like most other teleost fishes exhibit remarkably labile sex determination mechanisms (Munday et al. 2006a; Kuwamura et al. 2007; Kiewek-Martínez et al. 2010; Avise 2011), where the timing of sexual differentiation is an important driver of variation (Warner 1984; Kazancıoğlu and Alonzo 2010). Despite this flexibility in sex determination and the existence of opportunities throughout their evolutionary history to exercise it, labrids hardly do so in favor of pathways alternative to sex-changed males.

Our results confirm the broad association between sex allocation and mating system (Erisman et al. 2009, 2013; Kazancıoğlu and Alonzo 2010) and provide some of the first quantitative comparative evidence to support predicted correlations between character states (Robertson and Choat 1974; Choat and Robertson 1975; Robertson and Warner 1978; Warner and Robertson 1978; Warner 1984). The nature and timing of transitions we estimated on the MCC tree do not clearly resolve which of the two sets of traits might be driving change in the other. However, results of the evolutionary correlation analyses show a greater effect of mating system on sex allocation than the reverse (i.e. 3/4 transition rates and/or probabilities are dependent on mating system and 2/4 are dependent on sex allocation; fig. 6). Collectively, our results demonstrate that mating behavior with varying degrees of male size advantage can induce evolutionary change in a complex life-history trait.

The size-advantage model predicts that strong social control over females will provide males with the greatest size advantage, thus enabling the strongest selection for protogyny (Robertson and Choat 1974; Warner 1975, 1984; Emlen and Oring 1977; Robertson and Warner 1978; Warner and Robertson 1978). In haremic polygyny, dominant males are able to control the mating and sex change of subordinate females (Robertson and Choat 1974; Nemtzov 1985; Lutnesky 1994; Morrey et al. 2002). Because of this strict social control, non-dominant, primary males are evolutionarily unfit – they are unable to gain access to females for reproduction – and haremic species are predicted to exhibit monandric protogyny (Robertson and Choat 1974). This line of reasoning explains how the expression of haremic polygyny is able to influence or change sex allocation to monandric protogyny. Indeed, we recovered this pattern of character change for the clade containing *Coris julis* and its close congeneric relatives (fig. 2 – see node 8 on the mating system character map and node 10 on the sex allocation character map). In contrast, at the base of the *Sparisoma* clade it appears that monandry was in place prior to transitions to haremic polygyny (fig. 2). Furthermore, Bayesian analyses show the rate and probability of transitions to monandry are not dependent on the type of polygynous mating (fig. 6*B*). This suggests that monandric protogyny may also be a sustainable sex allocation strategy under lower male-size advantage characteristic of lek-like mating. Such a character combination may arise if sex-changed males are able to limit the reproductive success of other males without restricting female movement to the same degree as in haremic systems. For example, pronounced visual traits like colour pattern or display behavior combined with unyielding female mate choice may reduce the chances of primary males gaining access to mates to near zero.

Changes in the distribution of resources, or other environmental factors can limit the ability of males to monopolize females, resulting in the breakdown of haremic polygyny (Emlen and Oring 1977). When this happens differential dominance relationships between males are expected to form, as exist in lek-like polygyny (Emlen and Oring 1977). This notion is supported by the high rate of reversal we estimated from haremic to lek-like polygyny (fig. 3, 5*A*). Transitions of this nature would create the potential for primary males to gain reproductive access to mates and establish evolutionary fitness, thereby facilitating transitions to diandric protogyny (Robertson and Choat 1974; Emlen and Oring 1977; Robertson and Warner 1978; Warner and Robertson 1978). Several clades show this pattern where lineages that first expressed lek-like polygyny transition to diandric protogyny (fig. 2 – see nodes 1, 4 and 6 of the sex allocation character map). However, other species exhibit the opposite order of character change, namely *Scarus rivulatus*, *Scarus globiceps*, *Scarus ferrugineus*, and *Scarus forsteni* (fig. 2). Bayesian analyses show that transitions to diandry have the greatest state-dependence on whether lineages express lek-like or haremic mating, with a rate coefficient of zero assigned to haremic lineages 79% of the time (fig. 6*B*). Taken together, our results suggest that labrids rarely decrease their expression of protogyny, especially when their mating behavior permits strong male size advantage.

Gonochorism is predicted to evolve when males and females have similar size-specific reproductive expectations (Warner 1975). When the density of males increases due to local population growth, the ability of small males to gain reproductive access to females also increases and the dilution of gametes reduces the reproductive advantage of large males (Warner 1984; Muñoz and Warner 2003, 2004). In other words, as mating becomes more promiscuous selection for protogyny decreases (Robertson and Warner 1978; Warner 1984; Hoffman 1985). Transitions from protogyny to gonochorism have been associated with transitions to promiscuous mating systems, such as group spawning for several groups of teleost fishes including labrids (Erisman et al. 2009, 2013). Our results also support this pattern of character change and show that, for this combination of traits, transitions to gonochorism have the greatest state-dependence on whether lineages express promiscuity or not, with a rate coefficient of zero assigned to polygynous lineages 69% of the time (fig. 6*A*).

We also find support for the main prediction of the SAM, that as male size-advantage increases (i.e. the rate of increase in reproductive value with size becomes greater in males), selection for protogynous sex change increases. However, this pattern was not apparent when we restricted the analyses to species with specific types of polygyny and protogyny. This suggests that once evolved, the extent to which protogynous sex change is expressed is highly adaptable. Labrids have long been a model system for studying the adaptive significance of sequential hermaphroditism (Darwin 1871; Ghiselin 1969; Robertson 1972; Robertson and Choat 1974; Warner 1975; Warner et al. 1975; Leigh et al. 1976; Warner and Robertson 1978; Warner 1984; Muñoz and Warner 2004). Our description of the nature and timing of transitions in mating system and sex allocation builds upon considerable past research efforts, but remains dependent on the taxa included. Some of the patterns we recovered are consistent with previous ideas about the drivers of change in these complex traits and our results provide some of the first quantitative evidence for fishes showing that specific types of polygynous mating and protogynous sex change have evolved synergistically following predictions of the SAM. It is our hope that this work will spark new discussion about the interrelatedness of mating system and sex allocation and the factors capable of influencing them.

## Data availability

All molecular data are available from GenBank, accession numbers provided in table A1. Mating system and sex allocation data are provided in table A5. All tree files and datafiles are also deposited in Dryad (accession number pending publication).

## Acknowledgements

We wish to thank our colleagues who have spent considerable time and effort collecting detailed observational, demographic, life-history, and molecular data of wrasses and parrotfishes. Thanks to past and present members of the Wainwright lab, Ross Robertson and Howard Choat for stimulating discussion of the ideas and taxa involved; Bob Thomson, Michael Landis and Samantha Price for their support with the phylogenetic and comparative analyses; Yutong Song for assistance with searching the literature; and Tim Caro, Tamra Mendelson, Gail Patricelli, and Stephen Staples for their comments on previous iterations of this paper. J.R.H. was supported by the National Science Foundation’s Postdoctoral Research Fellowship in Biology for Research using Biological Collections (DBI-1523934) and parts of this work were supported by National Science Foundation Grants DEB-0717009 and DEB-1061981 to P.C.W.

## References

Adreani, M. S., B. E. Erisman, and R. R. Warner. 2004. Courtship and spawning behavior in the California sheephead, Semicossyphus pulcher (Pisces: Labridae). Environmental Biology of Fishes 71:13–19.

Aiello, B. R., M. W. Westneat, and M. E. Hale. 2017. Mechanosensation is evolutionarily tuned to locomotor mechanics. Proceedings of the National Academy of Sciences 114:4459–4464.

Alfaro, M., C. Brock, B. Banbury, and P. Wainwright. 2009. Does evolutionary innovation in pharyngeal jaws lead to rapid lineage diversification in labrid fishes? BMC Evolutionary Biology 9:255.

Anthes, N., H. Schulenburg, and N. K. Michiels. 2008. Evolutionary links between reproductive morphology, ecology, and mating behavior in opisthobranch gastropods. Evolution 62:900–916.

Avise, J. 2011. Hermaphroditism: A Primer on the Biology, Ecology, and Evolution of Dual Sexuality. Columbia University Press, New York.

Baeza, J. A., and R. T. Bauer. 2004. Experimental test of socially mediated sex change in a protandric simultaneous hermaphrodite, the marine shrimp Lysmata wurdemanni (Caridea: Hippolytidae). Behavioral Ecology and Sociobiology 55:544–550.

Baliga, V. B., and C. J. Law. 2016. Cleaners amongst wrasses: phylogenetics and evolutionary patterns of cleaning behavior within Labridae. Molecular Phylogenetics and Evolution 94:424–435.

Blomberg, S. P., T. Garland, and A. R. Ives. 2003.Testing for phylogenetic signal in comparative data: behavioral traits are more labile. Evolution 57:717–745.

Charnov, E. L. 1982. The Theory of Sex Allocation (Volume 18.). Princeton University Press, Princeton, NJ, NJ.

Choat, J. H., and D. R. Robertson. 1975. Protogynous hermaphroditism in fishes of the family Scaridae. Pages 263–283 in R. Reinboth, ed. Intersexuality in the Animal Kingdom. Springer-Verlag, Berlin.

Colin, P. L., and L. J. Bell. 1991. Aspects of the spawning of labrid and scarid fishes (Pisces: Labroidei) at Enewetak Atoll, Marshall Islands with notes on other families. Environmental Biology of Fishes 31:229–260.

Collin, R. 2006. Sex ratio, life history invariants, and patterns of sex change in a family of protandrous gastropods. Evolution 60:735–745.

Cowen, R. K. 1990. Sex change and life history patterns of the labrid, *Semicossyphus pulcher*, across an environmental gradient. Copeia 1990:787–795.

Cowman, P. F., and D. R. Bellwood. 2011. Coral reefs as drivers of cladogenesis: expanding coral reefs, cryptic extinction events, and the development of biodiversity hotspots. Journal of Evolutionary Biology 24:2543–2562.

Darwin, C. 1871. The Descent of Man, and Selection in Relation to Sex. John Murray, London.

Drummond, A. J., S. Y. W. Ho, M. J. Phillips, and A. Rambaut. 2006. Relaxed phylogenetics and dating with confidence. PLoS Biology 4:699–710.

Drummond, A. J., M. A. Suchard, D. Xie, and A. Rambaut. 2012. Bayesian phylogenetics with BEAUti and the BEAST 1.7. Molecular Biology and Evolution 29:1969–1973.

Emlen, S. T., and L. W. Oring. 1977. Ecology, sexual selection, and the evolution of mating systems. Science 197:215–223.

Erisman, B. E., M. T. Craig, and P. A. Hastings. 2009. A phylogenetic test of the size-advantage model: evolutionary changes in mating behavior influence the loss of sex change in a fish lineage. The American Naturalist 174:E83–E99.

Erisman, B. E., C. W. Petersen, P. A. Hastings, and R. R. Warner. 2013. Phylogenetic perspectives on the evolution of functional hermaphroditism in teleost fishes. Integrative and Comparative Biology 53:736–754.

Eschmeyer, W. N., R. Fricke, and R. van der Laan. 2018. Catalog of Fishes: Genera, Species, References. (W. N. Eschmeyer, R. Fricke, & R. van der Laan, eds.).

Gernhard, T. 2008. The conditioned reconstructed process. Journal of Theoretical Biology 253:769–778.

Ghiselin, M. T. 1969. The evolution of hermaphroditism among animals. The Quarterly Review of Biology 44:189–208.

Hodge, J. R., and D. R. Bellwood. 2015. On the relationship between species age and geographical range in reef fishes: are widespread species older than they seem? Global Ecology and Biogeography 24:495–505.

Hodge, J. R., F. Santini, and P. C. Wainwright. 2019. Data from: A closer look at the co-evolution of protogynous sex change and polygynous mating in wrasses and parrotfishes (Family: Labridae). American Naturalist, Dryad Digital Repository, URL pending publication.

Hoffman, S. G. 1985. Effects of size and sex on the social organization of reef-associated hogfishes, Bodianus spp. Environmental Biology of Fishes 14:185–197.

Kass, R. E., and A. E. Raftery. 1995. Bayes factors. Journal of the American Statistical Association 90:773–795.

Kazancıoğlu, E., and S. H. Alonzo. 2010. A comparative analysis of sex change in Labridae supports the size advantage hypothesis. Evolution 64:2254–2264.

Kiewek-Martínez, M., V. Gracia-López, and C. Rodríguez-Jaramillo. 2010. Evidence of sexual transition in Leopard Grouper (*Mycteroperca rosacea*) individuals held in captivity. Hidrobiológica 20:213–221.

Kuwamura, T., S. Suzuki, N. Tanaka, E. Ouchi, K. Karino, and Y. Nakashima. 2007. Sex change of primary males in a diandric labrid Halichoeres trimaculatus: coexistence of protandry and protogyny within a species. Journal of Fish Biology 70:1898–1906.

Leigh, E. G., E. L. Charnov, and R. R. Warner. 1976. Sex ratio, sex change, and natural selection. Proceedings of the National Academy of Sciences of the United States of America 73:3656–3660.

Lutnesky, M. M. F. 1994. Density-dependent protogynous sex change in territorial-haremic fishes: models and evidence. Behavioral Ecology 5:375–383.

Mank, J. E., D. E. L. Promislow, and J. C. Avise. 2006 Evolution of alternative sex-determining mechanisms in telesot fishes. Biological Journal of the Linnean Society 87:83–93.

McBride, R. S., and M. R. Johnson. 2007. Sexual development and reproductive seasonality of hogfish (Labridae: Lachnolaimus maximus), an hermaphroditic reef fish. Journal of Fish Biology 71:1270–1292.

McInerney, F. A., and S. L. Wing. 2011. The Paleocene–Eocene thermal maximum: A perturbation of carbon cycle, climate, and biosphere with implications for the future. Annual Review of Earth and Planetary Sciences 39:489–516.

Molloy, P. P., N. B. Goodwin, I. M. Côté, J. D. Reynolds, and M. J. G. Gage. 2007. Sperm competition and sex change: a comparative analysis across fishes. Evolution 61:640–652.

Morrey, C. E., Y. Nagahama, and E. G. Grau. 2002. Terminal phase males stimulate ovarian function and inhibit sex change in the protogynous wrasse Thalassoma duperrey. Zoological Science 19:103–109.

Munday, P. L., P. M. Buston, and R. R. Warner. 2006a. Diversity and flexibility of sex-change strategies in animals. Trends in Ecology & Evolution 21:89–95.

Munday, P. L., J. W. White, and R. R. Warner. 2006b. A social basis for the development of primary males in a sex-changing fish. Proceedings of the Royal Society B: Biological Sciences 273:2845–2851.

Muñoz, R. C., and R. R. Warner. 2003. A new version of the size-advantage hypothesis for sex change: incorporating sperm competition and size-fecundity skew. The American Naturalist 161:749–761.

Muñoz, R. C., and R. R. Warner. 2004. Testing a new version of the size-advantage hypothesis for sex change: sperm competition and size-skew effects in the bucktooth parrotfish, *Sparisoma radians*. Behavioral Ecology 15:129–136.

Nemtzov, S. C. 1985. Social control of sex change in the Red Sea razorfish Xyrichtys pentadactylus (Teleostei, Labridae). Environmental Biology of Fishes 14:199–211.

Payros, A., S. Ortiz, I. Millán, J. Arostegi, X. Orue-Etxebarria, and E. Apellaniz. 2015. Early Eocene climatic optimum: environmental impact on the North Iberian continental margin. Geological Society of America Bulletin 127:1632–1644.

Pagel, M., and A. Meade. 2006. Bayesian analysis of correlated evolution of discrete characters by reversible-jump Markov chain Monte Carlo. The American Naturalist 167:808–825.

Pagel, M., A. Meade, and D. Barker. 2004. Bayesian estimation of ancestral character states on phylogenies. Systematic Biology 53:673–684.

Paradis, E., J. Claude, and K. Strimmer. 2004. APE: Analyses of phylogenetics and evolution in R language. Bioinformatics 20:289–290.

Pitcher, T. J. 1992. The Behaviour of Teleost Fishes (2nd ed.). Springer Science & Business Media.

Policansky, D. 1982. Sex change in plants and animals. Annual Review of Ecology and Systematics 13:471–495.

Raftery, A. E. 1996. Hypothesis testing and model selection. Pages 163–188 in W. R. Gilks, S. Richardson, and D. J. Spiegelhalter, eds. Markov Chain Monte Carlo in Practice. Chapman & Hall, London.

Rambaut, A., M. Suchard, and A. J. Drummond. 2009. Tracer v1.5.

Renne, P. R., A. L. Deino, F. J. Hilgen, K. F. Kuiper, D. F. Mark, W. S. Mitchell, L. E. Morgan, et al. 2013. Time scales of critical events around the Cretaceous-Paleogene boundary. Science 339:684–687.

Robertson, D. R. 1972. Social control of sex reversal in a coral-reef fish. Science 177:1007–1009.

Robertson, D. R., and J. H. Choat. 1974. Protogynous hermaphroditism and social systems in labrid fish. Pages 217–224 in Proceedings of the Second International Coral Reef Symposium (Vol. 1). Brisbane, Australia.

Robertson, D. R., and R. R. Warner. 1978. Sexual patterns in the labroid fishes of the western Caribbean, II: the parrotfishes (Scaridae). Smithsonian Contributions to Zoology 255:1–26.

Robertson, R. D., R. Reinboth, and R. W. Bruce. 1982. Gonochorism, protogynous sex-change and spawning in three sparisomatinine parrotfishes from the western Indian Ocean. Bulletin of Marine Science 32:868–879.

Ronquist, F., and J. P. Huelsenbeck. 2003. MrBayes 3: Bayesian phylogenetic inference under mixed models. Bioinformatics 19:1572–1574.

Ross, R. M. 1990. The evolution of sex-change mechanisms in fishes. Environmental Biology of Fishes 29:81–93.

Sadovy de Mitcheson, Y., and M. Liu. 2008. Functional hermaphroditism in teleosts. FISH and FISHERIES 9:1–43.

Sadovy, Y., and D. Y. Shapiro. 1987. Criteria for the diagnosis of hermaphroditism in fishes. Copeia 1987:136–156.

Santini, F., L. Sorenson, and M. E. Alfaro. 2016. Phylogeny and biogeography of hogfishes and allies (*Bodianus*, Labridae). Molecular Phylogenetics and Evolution 99:1–6.

Shapiro, D. Y. 1987. Differentiation and evolution of sex change in fishes. BioScience 37:490–497.

Smith, C. L. 1975. The Evolution of Hermaphroditism in Fishes. Pages 295–310 in R. Reinboth, ed. Intersexuality in the Animal Kingdom. Springer Berlin Heidelberg, Berlin, Heidelberg.

Taborsky, M. 1998. Sperm competition in fish: ‘bourgeois’ males and parasitic spawning. Trends in Ecology & Evolution 13:222–227.

Todd, E. V, H. Liu, S. Muncaster, and N. J. Gemmell. 2016. Bending genders: The biology of natural sex change in fish. Sexual Development: Genetics, Molecular Biology, Evolution, Endocrinology, Embryology, and Pathology of Sex Determination and Differentiation 10:223–241.

Wainwright, P. C., F. Santini, D. R. Bellwood, D. R. Robertson, L. A. Rocha, and M. E. Alfaro. 2018. Phylogenetics and geography of speciation in New World *Halichoeres* wrasses. Molecular Phylogenetics and Evolution 121:35–45.

Warner, R. R. 1975. The adaptive significance of sequential hermaphroditism in animals. The American Naturalist 109:61–82.

Warner, R. R. 1982. Mating systems, sex change and sexual demography in the rainbow wrasse, *Thalassoma lucasanum*. Copeia 1982:653–661.

Warner, R. R. 1984. Mating behavior and hermaphroditism in coral reef fishes: the diverse forms of sexuality found among tropical marine fishes can be viewed as adaptations to their equally diverse mating systems. American Scientist 72:128–136.

Warner, R. R. 1988. Sex change and the size-advantage model. Trends in Ecology & Evolution 3:133–136.

Warner, R. R., D. L. Fitch, and J. D. Standish. 1996. Social control of sex change in the shelf limpet, Crepidula norrisiarum: size-specific responses to local group composition. Journal of Experimental Marine Biology and Ecology 204:155–167.

Warner, R. R., and S. G. Hoffman. 1980a. Local population size as a determinant of mating system and sexual composition in two tropical marine fishes (*Thalassoma* Spp.). Evolution 34:508–518.

Warner, R. R., and S. G. Hoffman. 1980b. Population density and the economics of territorial defense in a coral reef fish. Ecology 61:772–780.

Warner, R. R., and P. Lejeune. 1985. Sex change limited by paternal care: a test using four Mediterranean labrid fishes, genus Symphodus. Marine Biology 87:89–99.

Warner, R. R., and D. R. Robertson. 1978. Sexual patterns in the labroid fishes of the western Caribbean, I: the wrasses (Labridae). Smithsonian Contributions to Zoology 254:1–27.

Warner, R. R., D. R. Robertson, and E. G. Leigh Jr. 1975. Sex change and sexual selection. Science 190:633–638.

Westneat, M. W., and M. E. Alfaro. 2005. Phylogenetic relationships and evolutionary history of the reef fish family Labridae. Molecular Phylogenetics and Evolution 36:370–390.

Xie, W., P. O. Lewis, Y. Fan, L. Kuo, and M.-H. Chen. 2011. Improving marginal likelihood estimation for Bayesian phylogenetic model selection. Systematic Biology 60:150–160.

