## Supplementary material for "Correlated evolution of sex allocation and mating system in wrasses and parrotfishes": Tables A2, A4, A7

**This PDF file includes:**

Tables A2, A4, A7

Captions for Tables A1–A6

**Other Supplemental Material for this manuscript includes:**

Tables A1, A3, A5, A6 (Excel file)

**Table A1:** GenBank accession numbers of molecular sequences of wrasses and parrotfishes used in this study. Species designations are based on the Catalog of Fishes (Eschmeyer et al., 2018), accessed 20th March 2018. (–) indicates an unavailable sequence. (*) indicates sequences courtesy of G. Bernardi. AAN indicates sequences that are awaiting GenBank accession numbers.

(separate file)

**Table A2:** Molecular sampling coverage

| Gene region | No. species with data | % nominal species (n = 670) | % species in alignment (n = 403) | Mean |
|---|---|---|---|---|
| Mitochondrial loci: | | | | 65.57% |
| 12S | 238 | 35.52% | 59.06% | |
| 16S | 330 | 49.25% | 81.89% | |
| COI | 314 | 46.87% | 77.92% | |
| CytB | 175 | 26.12% | 43.42% | |
| Nuclear loci: | | | | 37.88% |
| RAG2 | 165 | 24.63% | 40.94% | |
| S7 | 128 | 19.10% | 31.76% | |
| TMO4c4 | 165 | 24.63% | 40.94% | |
| Mean | 216 | 32.30% | 53.70% | |

**Table A3:** Models of molecular evolution with the highest posterior probability. Models were sampled by the reversible-jump Markov chain Monte Carlo analysis across the entire general time reversible model space. Two independent runs were performed in MrBayes to evaluate convergence. For each partition, the model with the highest posterior probability between the two runs was used to specify corresponding parameters and priors in the BEAST analysis.
(separate file)

**Table A4:** Coral reef fish fossil data and other evidence used to temporally calibrate phylogenetic trees. Age constraints were used to convert the 50% majority-rule consensus tree into an ultrametric tree. Fossil calibrations were specified by placing exponential priors on nodes representing the most recent common ancestor (MRCA) of the specified genera. The offset refers to the minimum hard bound of the exponential prior distribution corresponding to the minimum age of the fossil; 95% refers to the soft upper bound of the exponential prior distribution and the mean describes its shape.

| | | Chronos age constraints | | BEAST fossil calibrations | | | |
|---|---|---|---|---|---|---|---|
| MRCA | Node No. | Min. age | Max. age | Offset | 95% | Mean | Evidence |
| *Kyphosus/Epinephelus* | 407 | 56 | 66 | – | – | – | legacy calibration |
| Labridae | 409 | 50 | 73.6 | 50 | 120 | 23.35 | *Phyllopharyngodon longipinnis* |
| *Tautoga/Tautogolabrus* | 412 | 15 | 24.3 | 15 | 50 | 11.7 | *Tautoga* sp. |
| *Symphodus* | 415 | 14 | 16.8 | 14 | 50 | 12.02 | *Symphodus westneati* |
| *Bolbometopon/Cetoscarus* | 514 | 5.5 | 17.7 | 5.3 | 50 | 14.92 | *Bolbometopon* sp. |
| *Calotomus/Sparisoma* | 516 | 20.9 | 35.5 | 14 | 50 | 12.02 | *Calotomus preisli* |
| *Achoerodus/Pseudodax* | 754 | 20 | 41 | 20 | 50 | 10.02 | *Trigondon jugleri* |

Sources. – (Carnevale and Godfrey n.d.; Bellwood 1990; Bellwood and Schultz 1991; Schultz and Bellwood 2004; Near et al. 2013; Carnevale 2015)

### References

Bellwood, D. R. 1990. A new fish fossil Phyllopharyngodon longipinnis gen. et sp. nov. (Family Labridae) from the Eocene, Monte Bolca, Italy. Studi e Ricerche sui Giacimenti Terziari di Bolca, Museo Civico di Storia Naturale. Verona 6:149–160.

Bellwood, D. R., and O. Schultz. 1991. A review of the fossil record of the parrotfishes (Labroidei: Scaridae) with a description of a new *Calotomus* species from the middle Miocene (Badenian) of Austria. Annalen Naturhistorisches Museum Wien 92A:55–71.

Carnevale, G. 2015. Middle Miocene wrasses (Teleostei, Labridae) from St. Margarethen (Burgenland, Austria). Palaeontographica. Abt. A: Palaeozoology - Stratigraphy 304:121–159.

Carnevale, G., and S. J. Godfrey. n.d. Miocene bony fishes of the Calvert, Choptank, St. Marys and Eastover Formations, Chesapeake Group, Maryland and Virginia. Smithsonian Contributions to Paleobiologylogy.

Near, T. J., A. Dornburg, R. I. Eytan, B. P. Keck, W. L. Smith, K. L. Kuhn, J. A. Moore, et al. 2013. Phylogeny and tempo of diversification in the superradiation of spiny-rayed fishes. Proceedings of the National Academy of Sciences USA 110:12738–12743.

Schultz, O., and D. R. Bellwood. 2004. *Trigonodon oweni* and *Asima jugleri* are different parts of the same species *Trigonodon jugleri*, a chiseltooth wrasse from the lower and middle Miocene in Central Europe (Osteichthyes, Labridae, Trigonodontinae). Annalen Naturhistorisches Museum Wien 105A:287–305.

**Table A5:** Data on mating system and sex allocation by species

(separate file)

**Table A6:** The total number of extant, nominal species for each labrid genus; the number included in the phylogenetic analysis, and the resultant percentage of species sampled. Species descriptions were obtained from the Catalog of Fishes (Eschmeyer et al., 2018), accessed 20th March 2018.

(separate file)

**Table A7:** Log marginal likelihoods estimated from dependent and independent models of character evolution, and corresponding log Bayes Factors. Three independent iterations were performed to test for correlated evolution between polygyny and protogyny (full dataset = 89 species), and between haremic polygyny and monandric protogyny (reduced dataset = 70). Log Bayes Factors indicate evidence against the null hypothesis (independent model), with values < 2 indicating weak evidence, 2–5 indicating positive evidence, and > 5 indicating strong evidence.

| Traits tested | Dependent model | Independent model | log Bayes Factor |
|---|---|---|---|
| Promiscuity & Polygyny | | | |
| Gonochorism & Protogyny: | | | |
| | –55.55 | –58.86 | 6.62 |
| | –56.16 | –58.75 | 5.17 |
| | –55.99 | –58.84 | 5.70 |
| Lek-like & Haremic Polygyny | | | |
| Diandric & Monandric Protogyny: | | | |
| | –80.59 | –82.19 | 3.20 |
| | –80.35 | –82.20 | 3.70 |
| | –80.39 | –82.22 | 3.65 |
